# Methylotrophic yeast *Candida boidinii* enhances the colonization of plant growth-promoting yeast *Papiliotrema laurentii* in the phyllosphere

**DOI:** 10.1101/2025.07.18.665470

**Authors:** Kana Shigeta, Kosuke Shiraishi, Moritz Schroll, Rebekka Lauer, Frank Keppler, Yasuyoshi Sakai, Hiroya Yurimoto

## Abstract

Methanol-utilizing microbes are ubiquitous in the phyllosphere, where they assimilate methanol released from pectin, the major component of the plant cell wall. While methylotrophic bacteria *Methylobacterium* spp. are well studied for their symbiotic relationships with the host plants, the ecology and functional roles of methylotrophic yeasts on plants remain poorly understood. In the effort to isolate yeasts from 26 phyllosphere samples, we identified *Candida boidinii* as the only methylotrophic yeast, while the remaining isolates, categorized into 17 species in 12 genera, lacked this metabolic trait. To obtain insight into the role of methylotrophic yeasts in the phyllosphere, we investigated the interaction of *C. boidinii* with a plant growth-promoting yeast (PGPY), *Papiliotrema laurentii*, one of the identified yeast species during isolation. We found that the colonization of *P. laurentii* was enhanced by the presence of *C. boidinii* on *Arabidopsis thaliana* leaves. Co-cultivation assays revealed that the cell yield of *P. laurentii* was enhanced by *C. boidinii* during cultivation on pectin and that the methanol-utilizing ability and pectin methylesterase (PME) activity of *C. boidinii* contributed to this enhancement. Stable carbon isotope labeling of pectin methylester groups unambiguously confirmed their assimilation by *C. boidinii*, but not by *P. laurentii*. These findings suggest that *C. boidinii* not only survives in the phyllosphere by utilizing pectin-derived methanol but also contributes to the fitness of other yeast species through metabolic cooperation. This study provides new insights into the niche construction and survival strategies of phyllosphere methylotrophic yeasts, highlighting their potential role in shaping microbial community dynamics and promoting beneficial plant-microbe interactions.

## Introduction

The phyllosphere, the aerial part of plants, represents one of the largest microbial habitats on Earth (Ruinen, 1956). The area of both sides of the plant leaves is estimated to be 6.4 × 10^8^ km^2^ (Morris and Kinkel, 2002), where a wide array of microbes reside. Among the phyllosphere microbes, bacterial populations are generally most abundant, reaching about 10^7^ colony-forming units (CFU) per gram of leaf on plates (Schlechter et al., 2019). However, other microbes also colonize plant leaves, including filamentous fungi ranging from 10^2^ to 10^8^ and yeasts ranging from 10 to 10^10^ per gram of leaf (Thompson et al., 1993, Inácio et al., 2002). Despite the enormous numbers of microbes, the phyllosphere is recognized as a challenging environment characterized by fluctuations in temperature, UV radiation, humidity and nutrient availability. To survive in such a harsh environment, microbial colonizers of the phyllosphere have evolved strategies to persist and interact with the host plant and neighboring microbes (Vorholt, 2012).

One critical aspect of such survival strategies in the phyllosphere is the acquisition of nutrients, in particular carbon sources. Methanol stands out as one of the most abundant organic compounds from plants, and is known to be utilized as a carbon source for a distinct type of microbes (Nemecek-Marshall et al., 1995, Fall and Benson, 1996). Methanol is released as a byproduct of plant cell wall remodeling, specifically through the hydrolysis of the methylester group of pectin by plant-derived pectin methylesterases (PMEs). Pectin comprises up to 35% of the primary cell wall, and it consists largely of homogalacturonan, a polymer of α-1,4-linked-galacturonic acid, some of which are methylesterified at the C-6 position (O’Neill et al., 1990, Ridley et al., 2001, Dorokhov et al., 2018).

The distinct group of microbes adapted to utilize methanol are methanol-utilizing microbes, organisms capable of utilizing methanol as their sole carbon and energy source. Among them, methylotrophic bacteria, members of the genus *Methylobacterium* spp., are known to be dominant inhabitants of the phyllosphere (Vorholt, 2012). While methylotrophic bacteria have been extensively studied, methylotrophic yeasts remain comparatively underexplored despite their frequent isolation from leaf surfaces (Santos et al., 2015, Nakase et al., 2010). These yeasts share the ability to utilize methanol but differ fundamentally in cellular organization and molecular mechanism of metabolic regulation from methylotrophic bacteria (Yurimoto et al., 2011). Our previous studies demonstrated that the methylotrophic yeast *Candida boidinii* proliferates on growing *A. thaliana* leaves by assimilating methanol as a carbon source, and regulates the gene expression of key metabolic enzymes such as alcohol oxidase (*AOD1*) and dihydroxyacetone synthase (*DAS1*) in response to diurnally fluctuating local methanol concentration (Kawaguchi et al., 2011). We also showed that available nitrogen sources for *C. boidinii* change from nitrate to methylamine during plant aging and that the yeast deploys the major protein degradation mechanism, autophagy, to adapt to such a change (Shiraishi et al., 2015). However, broader ecology and physiological roles of methylotrophic yeasts in the phyllosphere remain largely uncharacterized.

Some plant-associated microbes establish symbiotic relationships with the host plants. While such studies on bacteria are more advanced, as exemplified by methylotrophic bacteria, which induce growth promotion and contribute to stress tolerance (Koshy et al., 2025). Some yeast species have also been identified as plant growth-promoting yeasts (PGPYs), which have positive effects on plant growth, e.g., enhancing nutrient acquisition, modulating plant hormone levels and improving stress resilience (Nimsi et al., 2023). Such yeast species include *Rhodotorula mucilaginosa* (Ramos-Garza et al., 2016), *Papiliotrema laurentii* (previously reported as *Cryptococcus laurentii*) (Cloete et al., 2009) and *Candida tropicalis* (Amprayn et al., 2012). Despite these findings, little is known about how PGPYs interact with other microbes in the phyllosphere, including those with distinct metabolic capabilities such as methylotrophy.

In this study, we isolated and identified yeasts from the phyllosphere to clarify how methylotrophic yeasts contribute to microbial interactions on the leaf surface. Focusing on *C. boidinii*, the only yeast species capable of utilizing methanol among the isolates, we investigated its role in facilitating the colonization of a PGPY *P. laurentii*. Using the stable carbon isotope labeling technique, we unambiguously confirmed the assimilation of pectin methylester group by *C. boidinii*, but not by *P. laurentii*. Our findings suggest that pectin mediates yeast-yeast interspecies interaction in the phyllosphere, and that PME activity in *C. boidinii*, together with its ability to assimilate methanol, facilitates the colonization of *P. laurentii* through hydrolysis of the methylester group of pectin.

## Results

### Assimilation profile of pectin and methanol by yeasts isolated from the phyllosphere

Twenty-six phyllosphere samples were subjected to sequential enrichment culture in synthetic methanol (SM) liquid medium (Details of the yeast isolation are described in the Materials and Methods section). The cell suspensions obtained through enrichment culture were then spread onto SM agar plates. After 3-5 days, approximately 10 to 200 yeast-like colonies appeared on each plate, from which 3-8 colonies were isolated, resulting in a total of 170 isolates. Subsequently, all isolates were incubated in SM liquid medium in test tubes to determine their methanol utilization ability. After 3 days, 61 isolates showed substantial growth on methanol, whereas the other 109 isolates did not, suggesting that not only methylotrophic yeasts, but also non-methylotrophic yeasts, were obtained through our isolation.

Next, we performed sequence analysis of the D1/D2 domain in the 28S rDNA, and, if necessary, sequence analysis of the ITS regions and morphological observation, to identify the yeast species. As a result, all 61 isolates grown on methanol were identified as the methylotrophic yeast *C. boidinii*. Likewise, the other 109 isolates were identified and classified into one of the following 17 species in 12 genera; *Aureobasidium melanogenum, Cryptococcus* sp.*, Cystobasidium calyptogenae, Cystobasidium minutum, Cystobasidium slooffiae, Meyerozyma caribbica, Naganishia diffluens, Papiliotrema aurea, Papiliotrema laurentii, Pichia kluyveri, Pseudozyma hubeiensis, Pseudozyma pruni, Rhodosporidiobolus ruineniae, Rhodotorula paludigena, Rhodotorula toruloides, Sungouiella intermedia* and *Zalaria obscura*. The first isolate of each yeast species was designated as “representative yeast strain” in this study and used for further experiments (Supplemental Table 1).

Using the 18 representative yeast strains, we investigated their assimilation of methanol and pectin, the major source of methanol on plant leaves. After 3 days of incubation, only one strain, *C. boidinii* strain SML1, showed growth in SM liquid medium, while all 18 yeast strains grew in synthetic pectin (SP) liquid medium, with their final cell yield (OD_610_) ranging from 0.53±0.01 for *C. boidinii* strain SML1 to 2.49±0.20 for *A. melanogenum* strain GiL12 (Table 1).

**Table 1.** Final cell yield in SP and SM liquid medium of the representative yeast strains. The data are presented as the mean±standard error (SE; n = 3).

| Strain | Final cell yield (OD<sub>610</sub>) |  |
| --- | --- | --- |
| Medium | SP | SM |
| *Aureobasidium melanogenum* strain GiL12 | 2.49±0.20 | No growth |
| *Candida boidinii* strain SML1 | 0.53±0.01 | 2.00±0.08 |
| *Cryptococcus* sp. strain FCL26 | 1.59±0.16 | No growth |
| *Cystobasidium calyptogenae* strain PeF7 | 1.46±0.06 | No growth |
| *Cystobasidium minutum* strain GSMF2 | 1.90±0.07 | No growth |
| *Cystobasidium slooffiae* strain GiL13 | 1.80±0.13 | No growth |
| *Meyerozyma caribbica* strain GoKF3 | 0.83±0.11 | No growth |
| *Naganishia diffluens* strain ApF1 | 2.07±0.17 | No growth |
| *Papiliotrema aurea* strain FCL21 | 1.23±0.11 | No growth |
| *Papiliotrema laurentii* strain PeF4 | 1.35±0.06 | No growth |
| *Pichia kluyveri* strain GoKF5 | 1.25±0.26 | No growth |
| *Pseudozyma hubeiensis* strain SAL1 | 1.22±0.09 | No growth |
| *Pseudozyma pruni* strain FCL20 | 1.37±0.04 | No growth |
| *Rhodospiridiobolus ruineniae* strain GiL14 | 1.60±0.11 | No growth |
| *Rhodotorula paludigena* strain GSMF1 | 1.31±0.09 | No growth |
| *Rhodotorula toruloides* strain PeF8 | 1.65±0.12 | No growth |
| *Sungouiella intermedia* strain GoKF2 | 0.98±0.07 | No growth |
| *Zalaria obscura* strain GFF1 | 1.79±0.19 | No growth |

### *C. boidinii* enhances the colonization of *P. laurentii* on *A. thaliana* leaves

The isolation of *C. boidinii* as the only methylotrophic yeast from the tested plant samples implied that this yeast species is a common methylotrophic yeast living in the phyllosphere, interacting with other plant-associated microbes. Among the yeast species identified, we focused on *P. laurentii*. This yeast is known for its plant growth-promoting properties, such as the production of indole-3-acetic acid (IAA), siderophores, and solubilization of phosphorus and zinc (Kumla et al., 2020). We investigated whether *C. boidinii* and *P. laurentii* affect each other’s plant colonization ability.

First, we constructed the zeocin-resistant *C. boidinii* strain expressing Venus-fluorescent protein under the constitutive *ACT1* promoter, hereafter referred to as *C. boidinii* strain venus-zeo^R^. The laboratory strain, *C. boidinii* strain AOU-1, was used as a parental strain, since genomic information is available (Sakai and Tani, 1988). *A. thaliana* seeds were inoculated with *C. boidinii* strain venus-zeo^R^ (single inoculation) and then cultivated on Hoagland agar plates for two weeks. Confocal microscopy detected Venus-fluorescence around the stomata of *A. thaliana* leaves (Fig. 1A). We also performed a colony formation assay. The cell suspension obtained from the aerial part of *A. thaliana* was spread onto synthetic dextrose (SD) agar plates containing zeocin. After two days of incubation, Venus-fluorescent colonies appeared on the plate (Fig. 1B), and the average CFU per 1 mg of leaves was approximately 1,000 (Fig. 1C). These results suggest that *C. boidinii* inoculated on the seeds were transmitted to and colonized the phyllosphere of *A. thaliana*. Our previous research showed that *C. boidinii* strain *aod1Δ*, deficient in methanol utilization, cannot proliferate on *A. thaliana* leaves (Kawaguchi et al., 2011), indicating that methanol-utilizing ability is essential for leaf colonization. Therefore, we examined whether the methanol-utilizing ability of *C. boidinii* also contributes to its transmission from the seeds to leaves of *A. thaliana*. We inoculated a zeocin-resistant *C. boidinii* strain *aod1Δ* expressing Venus-fluorescent protein under the constitutive *ACT1* promoter, hereafter referred to as *C. boidinii* strain *aod1Δ* venus-zeo^R^, on seeds of *A. thaliana*. The result showed that *C. boidinii* strain *aod1Δ* venus-zeo^R^ did not form any colony on zeocin-containing SD plates (Fig. 1B-C), indicating that the methanol-utilizing ability is necessary for the migration of *C. boidinii* from the seeds to leaves and to colonize the phyllosphere of *A. thaliana*.

**Fig. 1.**
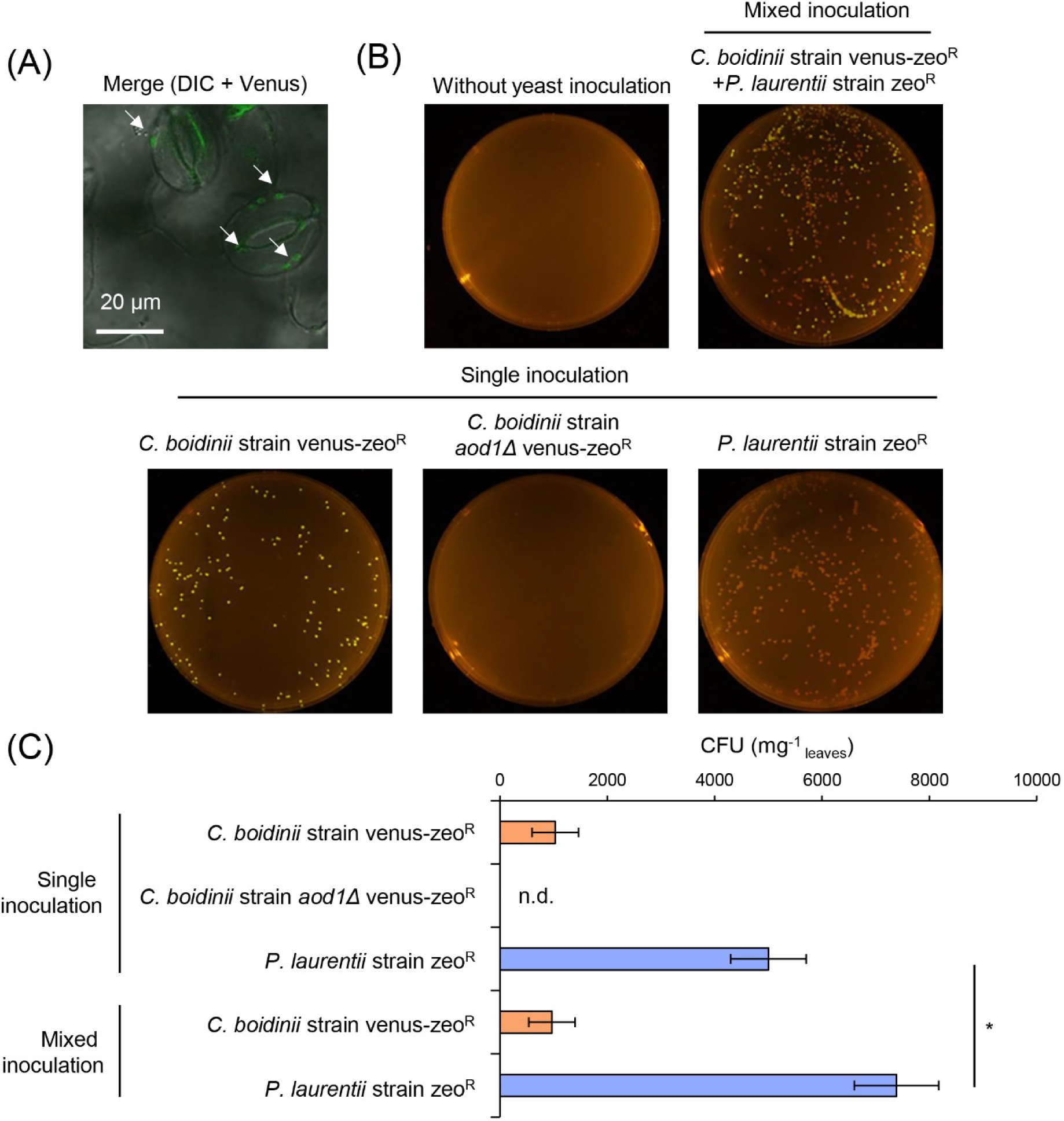
**Leaf colonization of *C. boidinii* and *P. laurentii* that were inoculated on seeds of *A. thaliana*.** (A) Microscopic observation of *C. boidinii* strain venus-zeo^R^ on the leaf surface of *A. thaliana*. White arrows represent the cells of *C. boidinii* strain venus-zeo^R^, Bar, 20 μm.(B-C) CFU of *C. boidinii* strain venus-zeo^R^ or *C. boidinii* strain *aod1Δ* venus-zeo^R^ and *P. laurentii* strain zeo^R^ on SD agar plate containing 150 µg/mL zeocin, (B) Images of the agar plates with the colonies of *C. boidinii* strain venus-zeo^R^, *C. boidinii* strain *aod1Δ* venus-zeo^R^ and *P. laurentii* strain zeo^R^, Quantification of (C) *C. boidinii* strain venus-zeo^R^, *C. boidinii* strain *aod1Δ* venus-zeo^R^ and *P. laurentii* strain zeo^R^ colonies appeared on the agar plates (CFU per mg of leaf). The data are presented as the mean±standard error (SE; n = 9). Significant differences are marked by *(*p* < 0.05; Student’s *t*-test). n.d., not detected

Similarly, we constructed the zeocin-resistant *P. laurentii* strain, hereafter referred to as *P. laurentii* strain zeo^R^, and performed a colony formation assay. *P. laurentii* strain PeF4, isolated from a leaf of a flowering cherry, was used as a parental strain (Supplemental Table 1). We observed a substantial number of colonies of *P. laurentii* strain zeo^R^ appeared on the tested plates (Fig. 1B), and the average CFU was around 5,000 (Fig. 1C). Given that no colony appeared from plant samples without yeast inoculation (Fig. 1B, our results suggested that *P. laurentii* inoculated on the seeds were transmitted to and colonized the phyllosphere of *A. thaliana*, similarly to *C. boidinii*.

Subsequently, *C. boidinii* strain venus-zeo^R^ and *P. laurentii* strain zeo^R^ were inoculated together (mixed inoculation) onto *A. thaliana* seeds. Two weeks after seeding, the aerial part of the plants was harvested for the colony formation assay. On the tested plates, colonies with and without Venus-fluorescence were observed (Fig. 1B), confirming the presence of both *C. boidinii* and *P. laurentii* in the phyllosphere of *A. thaliana* leaves. Quantification analysis revealed that an average of around 1,000 CFU of *C. boidinii* strain venus-zeo^R^ and 7,400 CFU of *P. laurentii* strain zeo^R^ were present (Fig. 1C). Interestingly, CFU of *P. laurentii* strain zeo^R^ under mixed inoculation conditions was significantly larger than that under single inoculation conditions, while CFU of *C. boidinii* strain venus-zeo^R^ did not show any significant difference between these two inoculation conditions (Fig. 1C). These results suggested that leaf colonization of *P. laurentii* is enhanced by the co-presence of *C. boidinii* in the phyllosphere of *A. thaliana*.

### *C. boidinii* enhances the cell yield of *P. laurentii* during co-cultivation on pectin

To explore the rationale behind the enhanced leaf colonization of *P. laurentii* by *C. boidinii*, we focused on their potential metabolic interaction involving pectin, a major component of the plant cell wall and one of the presumed carbon sources for phyllosphere yeasts (Blanco et al., 1999). Our previous study has suggested that *C. boidinii* produces and utilizes methanol derived from pectin (Nakagawa et al., 2000). To unambiguously confirm the pectin-utilizing capability of *C. boidinii*, we employed stable carbon isotopic labeling and analysis.

Polygalacturonic acid (PGA), a polymer of α-1,4-linked-galacturonic acid, was methylesterified at the C-6 position using ^13^C-labeled methanol to generate ^13^C-labeled methylesterified PGA. In principle, after hydrolysis of ^13^C-labeled methylester group of pectin, ^13^C-enriched methanol is produced and then converted to ^13^C-enriched CO_2_, which can be detected as the final product of this process by gas chromatography-isotope-ratio-mass spectrometry (Supplementary Fig. 2). As expected, a substantial amount of ^13^C-enriched CO_2_ was detected, when *C. boidinii* strain AOU-1 was cultivated in a synthetic liquid medium supplemented with ^13^C-labeled methylesterified PGA for 24 h. In contrast, cultivation in a synthetic liquid medium supplemented with unlabeled methylesterified PGA did not result in any detectable change in δ^13^C-CO_2_ values (Fig. 2A). These results confirmed that *C. boidinii* strain AOU-1 assimilates the methylester group of pectin and converts it to CO_2_. In comparison, no significant increase in δ^13^C-CO_2_ values was observed for *P. laurentii* strain PeF4 under the same cultivation conditions (Fig. 2A), confirming that *P. laurentii* strain PeF4 is unable to utilize the methylester group of pectin. Considering that *C. boidinii* uses methanol as a carbon source in the phyllosphere (Kawaguchi et al., 2011), but *P. laurentii* does not (Table 1), we hypothesized that, by taking advantage of its methanol-utilizing ability, *C. boidinii* alters the structure of pectin through remodeling and degradation in a way that facilitates the pectin utilization of *P. laurentii*.

**Fig. 2.**
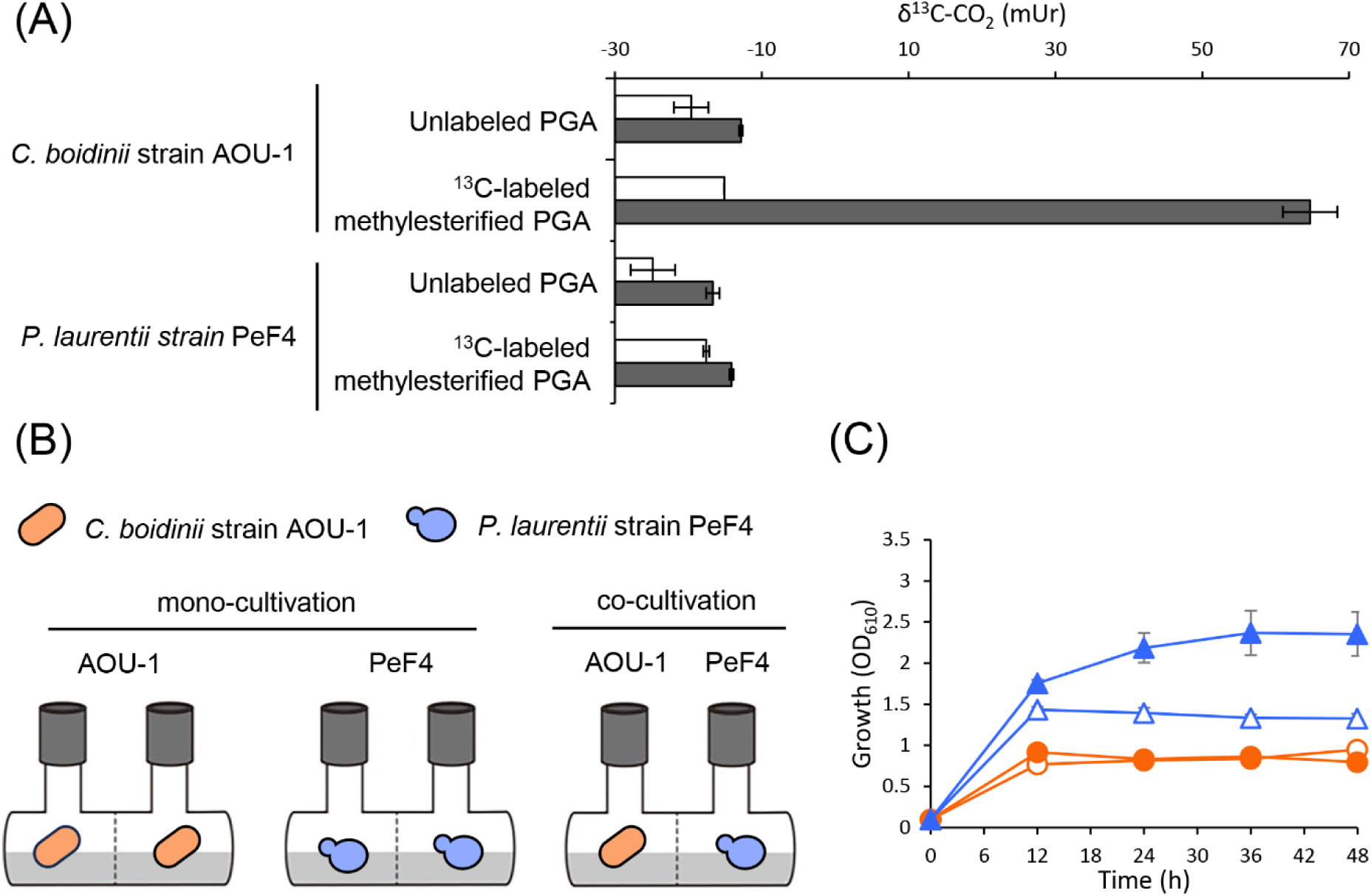
**pectin-utilizing capability of *C. boidinii* strain AOU-1 and *P. laurentii* strain PeF4 and the proliferation of *P. laurentii* strain PeF4 during co-cultivation with *C. boidinii* strain AOU-1 in SP liquid media.** (A) Utilization of methylester group of pectin by *C. boidinii* strain AOU-1 and *P. laurentii* strain PeF4. These two strains were cultivated in a synthetic liquid medium supplemented with unlabeled methylesterified PGA, and ^13^C-labeled methylesterified PGA, and stable carbon isotope values of CO_2_ (δ^13^C-CO_2_) was measured as the final product. An increase δ^13^C-CO_2_ value indicates enrichment of the heavier isotope ^13^C. Symbols: (white bars) start of cultivation; (gray bars) 24 h after cultivation. The data are presented as the mean±standard error (SE; n = 1-3). (B-C) Cultivation of *P. laurentii* strain PeF4 with *C. boidinii* strain AOU-1 in SP liquid media using *Beppu* flasks. (B) Schematic image of the cultivation method, using *Beppu* flasks. The images represent the mono- and co-cultivation of *C. boidinii* strain AOU-1 and *P. laurentii* strain PeF4. (C) Growth curve of *C. boidinii* strain AOU-1 and *P. laurentii* strain PeF4 during mono- and co-cultivation. Symbols: (empty circles) *C. boidinii* strain AOU-1 mono-cultivation; (filled circles) *C. boidinii* strain AOU-1 co-cultivation; (empty triangle) *P. laurentii* strain PeF4 mono-cultivation; (filled triangle) *P. laurentii* strain PeF4 co-cultivation. The data are presented as the mean±standard error (SE; n = 3).

To test our hypothesis, we first investigated the interaction between *C. boidinii* and *P. laurentii* in SP liquid medium using *Beppu* flasks. *Beppu* flasks consist of two rooms separated by a membrane filter that allows medium components, including pectin and its degradation products, to pass through and be maintained at the same level, but does not allow yeast cells to be mixed. *C. boidinii* strain AOU-1 and *P. laurentii* strain PeF4 were either mono-cultivated or co-cultivated in SP liquid medium (Fig. 2B). Under mono-cultivation conditions, the same yeast strains were incubated on both sides of the flask, and the cell growth was demonstrated as an average OD_610_ of both sides. No significant difference was observed in the growth curve of *C. boidinii* strain AOU-1 between mono- and co-cultivation conditions. In contrast, the cell yield of *P. laurentii* strain PeF4 was significantly enhanced when co-cultivated with *C. boidinii* strain AOU-1 (Fig. 2C). Cell yield of *P. laurentii* strain PeF4 under mono-cultivation peaked at 12 h after start of the incubation with OD_610_ 1.43±0.04, followed by a gradual decline, whereas under co-cultivation, it reached the maximum at 36 h with OD_610_ 2.37±0.27 and maintained nearly the same level even at 48 h. We calculated that the maximum cell yield during co-cultivation with *C. boidinii* strain AOU-1 was higher by 1.65 folds than that during mono-cultivation (Table 2).

**Table 2.**
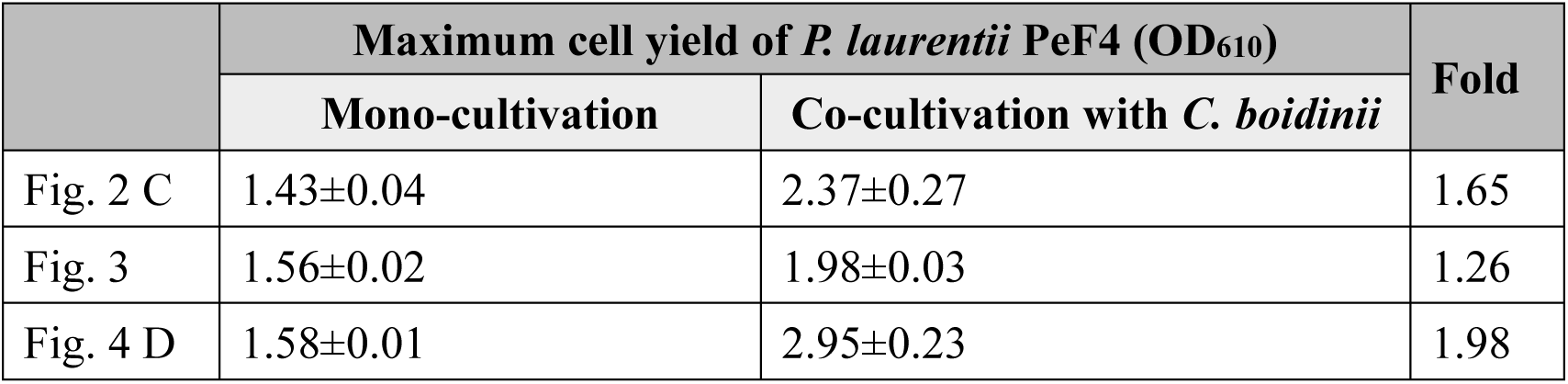
Comparison of the maximum cell yield of *P. laurentii* strain PeF4 during co-cultivation in SP media with *C. boidinii* strain AOU-1, *aod1Δ* or SML1 and that of *P. laurentii* strain PeF4 during mono-cultivation using *Beppu* flasks. Values of the maximum cell yield of *P. laurentii* strain PeF4 when co-cultivated with the respective *C. boidinii* strains relative to values of the maximum cell yield of *P. laurentii* strain PeF4 in mono-cultivation are shown as “Fold”. The data are presented as the mean±standard error (SE; n = 3).

|  | Maximum cell yield of *P. laurentii* PeF4 (OD<sub>610</sub>) |  | Fold |
| --- | --- | --- | --- |
|  | Mono-cultivation | Co-cultivation with *C. boidinii* |  |
| Fig. 2 C | 1.43±0.04 | 2.37±0.27 | 1.65 |
| Fig. 3 | 1.56±0.02 | 1.98±0.03 | 1.26 |
| Fig. 4 D | 1.58±0.01 | 2.95±0.23 | 1.98 |

Next, we investigated whether the cell yield of other representative yeast strains is also enhanced by co-cultivation with *C. boidinii* strain AOU-1 in SP liquid media. Similarly to *P. laurentii* strain PeF4, the cell yield of the other 16 representative yeast strains was enhanced when co-cultivated with *C. boidinii* strain AOU-1, with the degree ranging from 1.28 folds for *C. minutum* strain GSMF2 to 2.29 folds for *S. intermedia* strain GoKF2 (Supplemental Fig. 1A-P). We also performed co-cultivation of *P. laurentii* strain PeF4 with *R. ruineniae* strain GiL14 and *R. toruloides* strain PeF8 in SP liquid media to examine whether the cell yield of *P. laurentii* strain PeF4 is enhanced by other yeast species. However, there was no significant difference in the growth curve of *P. laurentii* strain PeF4, as well as *R. ruineniae* strain GiL14 and *R. toruloides* strain PeF8 across mono- and co-cultivation conditions (Supplemental Fig. 2A-B). Furthermore, *P. laurentii* strain PeF4 was co-cultivated with *C. boidinii* strain AOU-1 in SD liquid medium to test whether the enhancement of cell yield is observed. In contrast to SP liquid medium, the cell yield of *P. laurentii* strain PeF4 was not significantly affected by co-cultivation with *C. boidinii* strain AOU-1 in SD liquid medium (Supplemental Fig. 3). These results indicated that *C. boidinii* enhances the cell yield of the co-cultivated yeasts specifically during pectin utilization.

### Methanol utilization ability of *C. boidinii* contributes to enhancing the cell yield of *P. laurentii* during co-cultivation on pectin

Next, to determine whether the methanol-utilizing ability of *C. boidinii* contributes to the enhanced cell yield of other yeasts during co-cultivation, we used the *C. boidinii* strain *aod1Δ* for co-cultivation with *P. laurentii* strain PeF4 in SP liquid media. Similarly to *C. boidinii* strain AOU-1, the cell yield of *P. laurentii* strain PeF4 was enhanced when co-cultivated with *C. boidinii* strain *aod1Δ*, while no significant difference was observed in the growth curve of *C. boidinii* strain *aod1Δ* between mono- and co-cultivation conditions. Under mono-cultivation conditions, *P. laurentii* strain PeF4 showed the maximal cell yield at 12 h after start of the cultivation with OD_610_ 1.56±0.02, followed by a moderate decline. During co-cultivation with *C. boidinii* strain *aod1Δ*, the cell yield of *P. laurentii* strain PeF4 reached its peak at 36 h with OD_610_ 1.98±0.03 (Fig. 3), leading us to calculate that the maximum cell yield during co-cultivation with *C. boidinii* strain *aod1Δ* was higher by 1.26 folds than that during mono-cultivation (Table 2). Interestingly, the degree of cell yield enhancement by co-cultivation with *C. boidinii* strain *aod1Δ* was less than that by co-cultivated with *C. boidinii* strain AOU-1 (Table 2), suggesting that the methanol-utilizing ability of *C. boidinii* is important for the enhancement of the cell yield of *P. laurentii.* Subsequently, we investigated whether the presence of methanol derived from pectin, which *C. boidinii* strain *aod1Δ* cannot assimilate, negatively affects the cell growth of *P. laurentii* strain PeF4. *P. laurentii* strain PeF4 was cultivated in SP or synthetic pectin and methanol (SPM) media. The growth of *P. laurentii* strain PeF4 was not significantly different in SP from SPM liquid media (Supplemental Fig. 4), indicating that the cell yield of *P. laurentii* strain PeF4 was not affected by the presence of methanol. These results suggested that the enhancement of *P. laurentii* cell yield during co-cultivation in SP liquid media is supported by methanol utilization activity of *C. boidinii*, rather than by the removal of methanol to relieve potential growth inhibition.

**Fig. 3.**
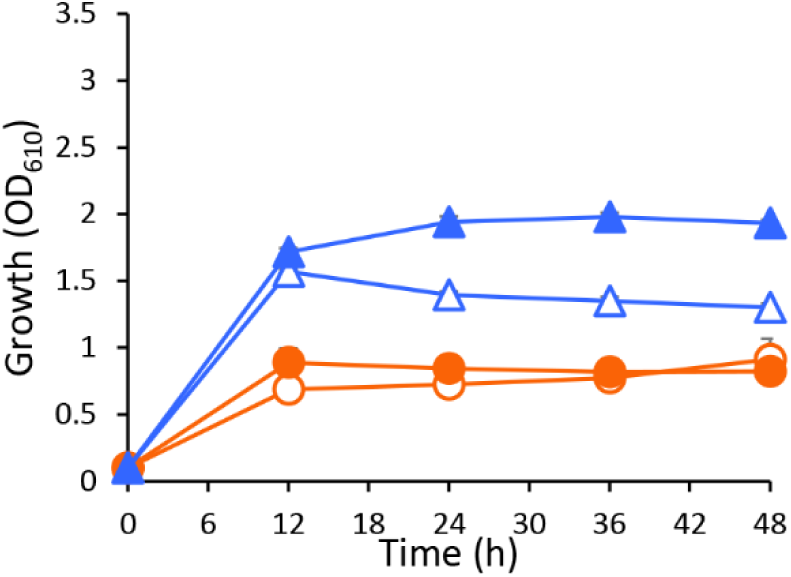
**The effect of *C. boidinii aod1* on the proliferation of *P. laurentii* strain PeF4 during co-cultivation in SP liquid media.** Growth curve of *C. boidinii* strain *aod1Δ* and *P. laurentii* strain PeF4 during mono- and co-cultivation in SP liquid media using *Beppu* flasks. Symbols: (empty circles) *C. boidinii* strain *aod1Δ* mono-cultivation; (filled circles) *C. boidinii* strain *aod1Δ* co-cultivation; (empty triangle) *P. laurentii* strain PeF4 mono-cultivation; (filled triangle) *P. laurentii* strain PeF4 co-cultivation. The data are presented as the mean±standard error (SE; n = 3).

### Pectin methylesterase (PME) activity of *C. boidinii* is one possible factor to enhance the cell yield of *P. laurentii* during co-cultivation on pectin

To gain further insights into the role of *C. boidinii*’s methanol utilization in the enhancement of *P. laurentii* cell yield, we examined the pectin methylesterase (PME) activity as it facilitates the methanol production from pectin. During cultivation in SP liquid medium, we measured the specific extracellular PME activity of *C. boidinii* strain AOU-1 and *P. laurentii* strain PeF4. PME activity was detected in *C. boidinii* strain AOU-1, but not in *P. laurentii* strain PeF4 (Fig. 4A), suggesting that *C. boidinii* hydrolyzes the methylester group of pectin, thereby modifying its structure in a way that promotes the assimilation of pectin by *P. laurentii* strain PeF4 and by other yeasts.

**Fig. 4.**
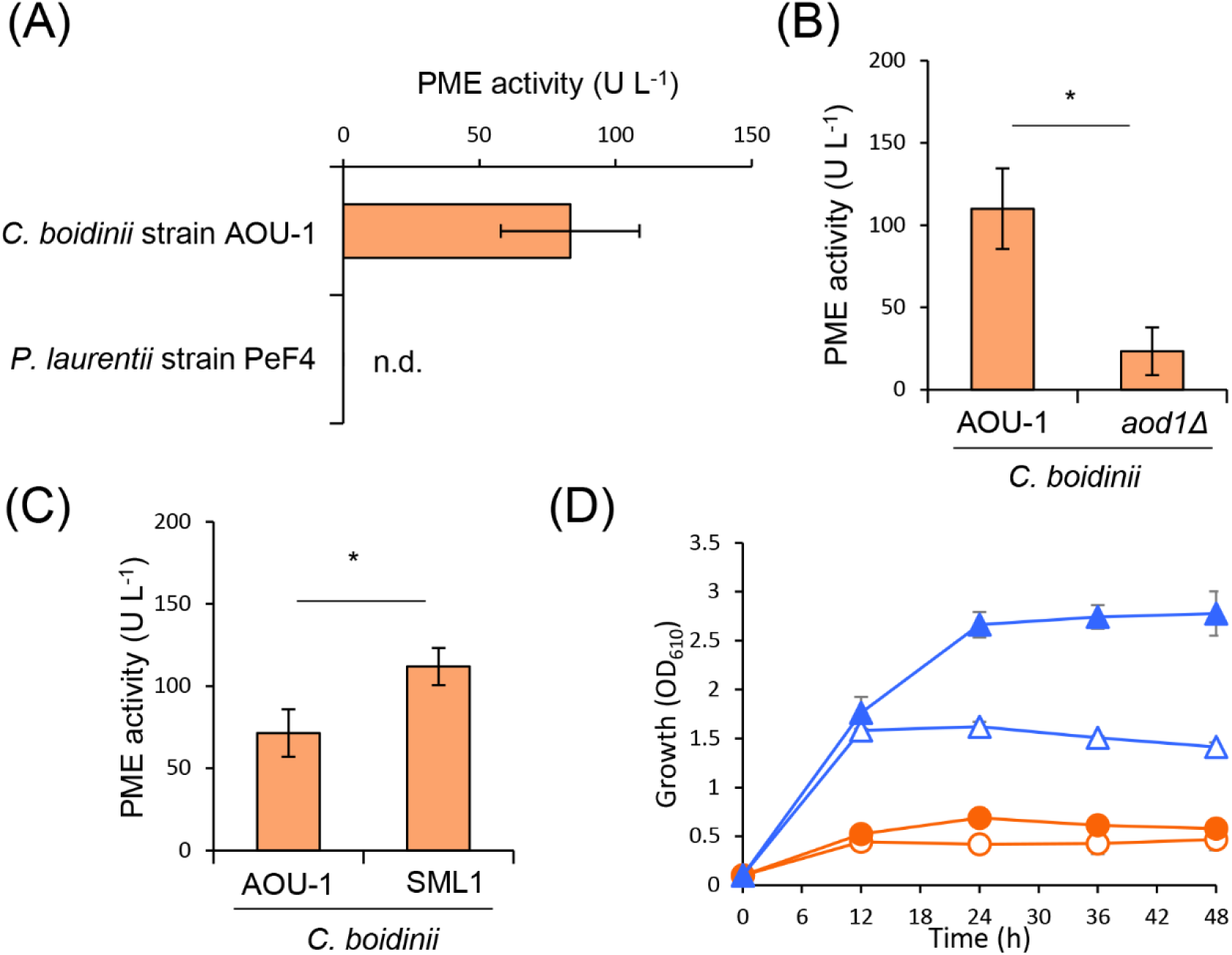
**The effect of *C. boidinii* PME activity on the proliferation of *P. laurentii* strain PeF4 during co-cultivation in SP liquid media.** Specific extracellular PME activity of (A) *C. boidinii* strain AOU-1 and *P. laurentii* strain PeF4, (B) *C. boidinii* strain AOU-1 and *C. boidinii* strain *aod1Δ,* (C) *C. boidinii* strain AOU-1 and *C. boidinii* strain SML1 during cultivation in SP liquid media. The data are presented as the mean±standard error (SE; n = 3 (A), 5 (B), 7 (C)) .n.d., not detected. Significant differences are marked by *(*p* < 0.05; Student’s *t*-test). (D) Growth curve of *C. boidinii* strain SML1 and *P. laurentii* strain PeF4 during mono- and co-cultivation in SP media using *Beppu* flasks. Symbols: (empty circles) *C. boidinii* strain SML1 mono-cultivation; (filled circles) *C. boidinii* strain SML1 co-cultivation; (empty triangle) *P. laurentii* strain PeF4 mono-cultivation; (filled triangle) *P. laurentii* strain PeF4 co-cultivation. The data are presented as the mean±standard error (SE; n = 3).

Next, PME activities of *C. boidinii* strain AOU-1 and *C. boidinii* strain *aod1Δ* were compared. We observed that the PME activity of *C. boidinii* strain *aod1Δ* was smaller than that of *C. boidinii* strain AOU-1 (Fig. 4B). Since *C. boidinii* strain *aod1Δ* is unable to assimilate methanol, we considered that methanol, produced from pectin, might suppress the PME activity. To test this, we compared the PME activity of *C. boidinii* strain *aod1Δ* during cultivation in SP and SPM liquid medium. The PME activity was significantly smaller in SPM liquid medium than in SP liquid medium (Supplemental Fig. 6), supporting our idea that PME activity is suppressed by the presence of methanol independently of its metabolism.

To explore whether higher PME activity correlates with greater ability to enhance the cell yield of other yeasts, we examined *C. boidinii* strain SML1, isolated from the phyllosphere. Notably, *C. boidinii* strain SML1 exhibited a higher PME activity than *C. boidinii* strain AOU-1 (Fig. 4C). We therefore conducted co-cultivation experiments using *C. boidinii* strain SML1 to assess whether enhanced PME activity contributes to improving the cell yield of *P. laurentii* strain PeF4. When *P. laurentii* strain PeF4 was mono-cultivated, it showed the maximal cell yield at 12 h with OD_610_ 1.58±0.01, while cell yield peaked at 48h with OD_610_ 2.95±0.23 during co-cultivation with *C. boidinii* strain SML1 (Fig. 4D). We calculated that the cell yield during co-cultivation with *C. boidinii* strain SML1 was higher by 1.98 folds than that during mono-cultivation, which was higher than co-cultivated with *C. boidinii* strain AOU-1 (Table 2). These results suggested that the PME activity of *C. boidinii*, supported by its methanol-utilizing ability, is a possible factor to enhance the cell yield of *P. laurentii* strain PeF4.

## Discussion

In this study, we demonstrated that the methylotrophic yeast *C. boidinii* enhances the colonization of *P. laurentii* in the phyllosphere of *A. thaliana*. One possible explanation, supported by stable carbon isotopic labeling and analysis, and co-cultivation assays, is the existence of a yeast-yeast interspecies interaction mediated by the utilization of pectin as a presumed carbon source in the phyllosphere. The PME activity of *C. boidinii*, along with its ability to assimilate methanol, contributes to this interaction by hydrolysis of the methylester group of pectin, thereby facilitating the pectin utilization by *P. laurentii* (Fig. 5).

**Fig. 5.**
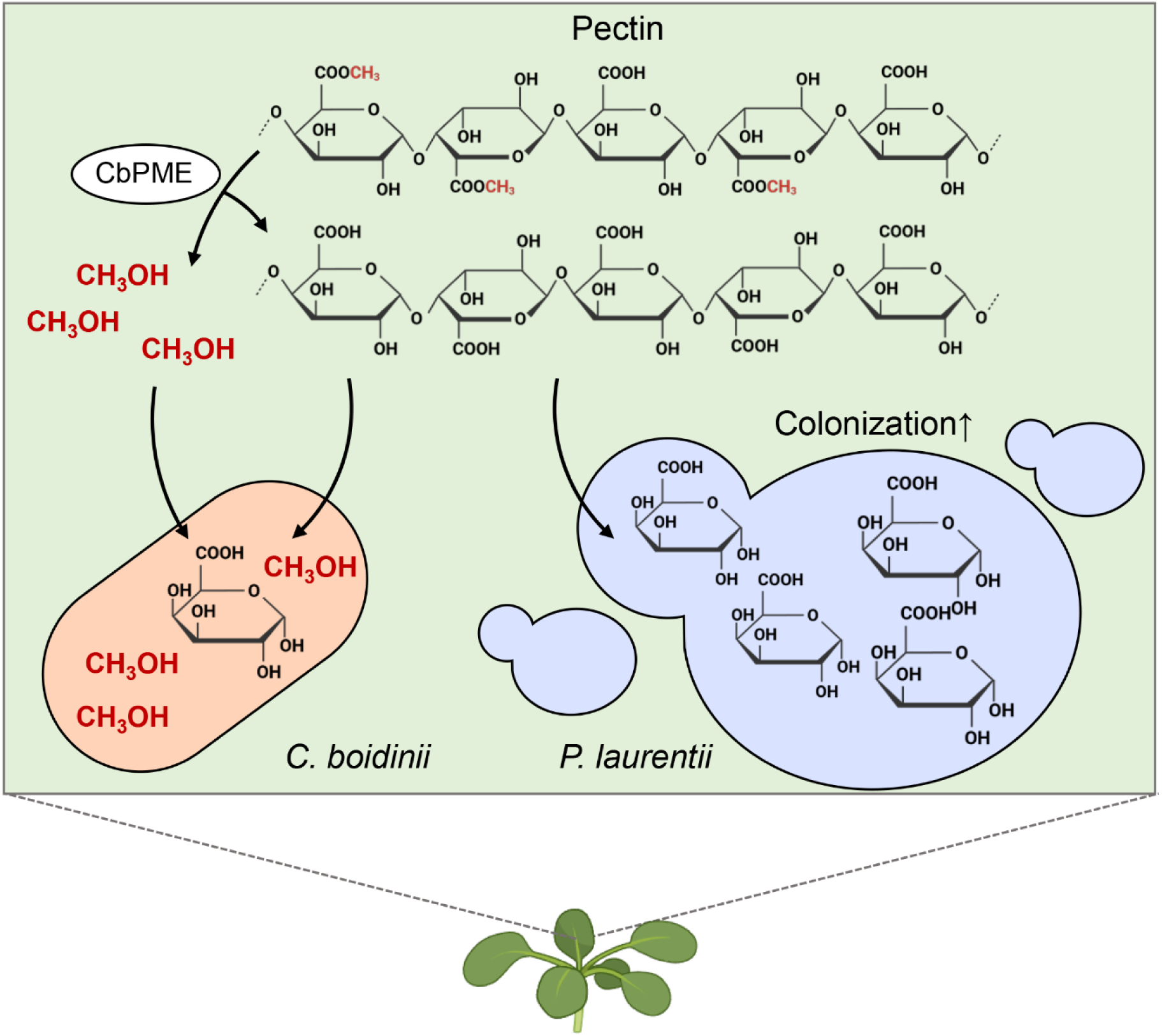
***Candida boidinii* enhances the colonization of *Papiliotrema laurentii* in the phyllosphere.** Methylotrophic yeast *C. boidinii* enhances the colonization of *P. laurentii* in the phyllosphere of *A. thaliana* possibly by facilitating the pectin utilization of *P. laurentii*. *C. boidinii* produces pectin methylesterase (PME), which hydrolyzes the methylester group of pectin, allowing *C. boidinii* to assimilate methanol and simultaneously providing *P. laurentii* with demethylesterified pectin, a more accessible carbon source, thereby promoting increased the leaf colonization of *P. laurentii*. Figure created with BioRender.com.

Our results suggested that *C. boidinii* produces and utilizes methanol derived from pectin (Fig. 2A and 4A), and in the phyllosphere, *C. boidinii* uses methanol as a carbon source (Kawaguchi et al., 2011). These results support a model in which *C. boidinii* hydrolyzes the methylester group of pectin via its PME activity and benefits from the released methanol as a carbon source, while the modified pectin becomes more accessible to *P. laurentii* and other phyllosphere yeasts (Fig. 2C and Supplemental Fig. 1A-P). While direct evidence for pectin leaching from plant leaves remains limited, previous studies suggest that pectin and its breakdown products are accessible to plant-associated microbes as carbon sources (Hoff et al., 2021, Kraut-Cohen et al., 2021). These results lead us to propose that *C. boidinii* facilitates the colonization of neighboring yeasts by providing metabolic benefits to others via pectin utilization.

Apart from the metabolic use, *C. boidinii* may alter the physical properties of pectin by its PME activity. Modification of pectin on plant surfaces has long been studied in the context of plant-pathogen interactions, where PME plays a key role in overcoming the cell wall barrier (Bellincampi et al., 2014). Unesterified pectin can form “egg-box” structures with Ca^2+^ that increase cell wall rigidity by cross-linking adjacent pectin chains (Wormit and Usadel, 2018). However, once the methylester group of pectin is hydrolyzed, it also becomes more chemically accessible and thus more susceptible to degradation by microbial enzymes such as pectate lyase (PL) and polygalacturonase (PG), leading to loosening of the cell wall structure (Wormit and Usadel, 2018). Recent studies have suggested that some symbiotic microbes, not just pathogens, secrete PME to facilitate colonization on plant surfaces (Su, 2023). Thus, PME activity may be a key functional trait shared by both pathogenic and symbiotic microbes to colonize leaf surfaces. Although our results did not directly demonstrate whether cell wall loosening promotes *P. laurentii* colonization, the observed yeast-yeast interspecies interaction in the phyllosphere may involve such structural modification. To investigate the molecular basis of pectin modification and assimilation by *C. boidinii*, we conducted a homology-based search for *PME* genes in *C. boidinii.* However, no clear candidates were identified, likely due to the poor conservation of *PME* genes among fungi (Markovic and Janecek, 2004). Out of the 18 yeast species we isolated in this study, only *P. hubeiensis* harbors a characterized *PME* gene (Konishi et al., 2013). *C. boidinii* was the only methylotrophic yeast among the phyllosphere isolates identified in this study. This species was originally isolated from soil and initially reported as *Kloeckera* sp. (Ogata et al., 1969). Since then, *C. boidinii* has been isolated from a wide range of environments, including soil, seawater, sap fluxes, fermentation sites and even hospital wards, suggesting that *C. boidinii* is a broadly distributed methylotrophic yeast in nature (Lachance et al., 2011).

Our previous work showed that *C. boidinii* proliferates on the leaf surface of *A. thaliana* (Kawaguchi et al., 2011). Building on this finding, the current study sheds light on its colonization strategy. We observed that *C. boidinii* is transmitted from seeds to leaves during early plant development, depending on its methanol utilization, enabling the yeast to establish residency on emerging leaf surfaces (Fig. 1A-D). Notably, *C. boidinii* cells were frequently observed near stomata (Fig. 1A). This spatial distribution pattern may be linked to methanol availability, as methanol is predominantly released from stomatal pores during plant metabolic activity (Fall and Benson, 1996). Recently, we reported that *Methylobacterium* sp. strain OR01, a methylotrophic bacterium, undergoes seed-to-leaf transmission and preferentially localizes near stomata. While the wild-type strain deploys methanol chemoreceptor- and flagellin protein-dependent methanol chemotaxis, methylotaxis, to actively reach stomatal cavities, even a non-motile mutant lacking flagella could colonize the phyllosphere (Katayama et al., 2025). Unlike motile bacteria, yeasts like *C. boidinii* lack flagella and are incapable of directed movement. Thus, their positioning around the stomata of the leaf surface may be governed not by active chemotaxis but likely by their passive accumulation in microenvironments, where methanol is more readily available to support growth.

Together, our findings show that *C. boidinii* plays a role in shaping microbial interactions in the phyllosphere. *C. boidinii* hydrolyzes plant-derived pectin through its PME activity, gaining access to methanol as a carbon source. At the same time, this modification makes pectin more accessible to other phyllosphere symbiotic yeasts, such as *P. laurentii*. Beyond its metabolic benefits, PME activity of *C. boidinii* may alter the physical properties of pectin in a way that facilitates microbial colonization. Future investigation into the molecular identity and regulation of *C. boidinii*’s PME activity will advance our understanding of yeast adaptation to the leaf surface environment and offer new strategies for engineering beneficial microbial consortia to support plant health.

## Materials and methods

### Medium

Yeast cells were grown at 28°C on the appropriate media described below. Yeast extract, peptone, and dextrose (YPD) medium consisted of 1% yeast extract, 2% bactopeptone, and 2% glucose. Synthetic dextrose (SD) medium was composed of 0.67% yeast nitrogen base without amino acids and 2% glucose. Synthetic pectin (SP) medium had 0.67% yeast nitrogen base without amino acids and 0.4% (w/v) pectin (DE, 70-75%; Nacalai Tesque). Synthetic methanol (SM) medium consisted of 0.67% yeast nitrogen base without amino acids and 0.5% (v/v) methanol. Synthetic liquid medium supplemented with ^13^C-labeled methylesterified PGA consisted of 0.67% yeast nitrogen base without amino acids and 0.5% (w/v) ^13^C-labeled PGA (1.8% methylester content, 5% ^13^C-labeled methylester group).

Synthetic liquid medium supplemented with unlabeled methylesterified PGA consisted of 0.67% yeast nitrogen base without amino acids and 0.5% (w/v) unlabeled methylesterified PGA (1.6% methylester content). Synthetic pectin and methanol (SPM) medium had 0.67% yeast nitrogen base without amino acids, 0.4% (w/v) pectin, and 0.7% (v/v) methanol.

*Escherichia coli* cells were grown at 37°C in LB medium (1% tryptone, 0.5% yeast extract, and 0.5% NaCl), supplemented with ampicillin (50 mg/L) when required. For the preparation of solid medium plates, the above media were supplemented with 2 % agar. All the SM media used for yeast isolation were supplemented with 1.5 mg/mL chloramphenicol to eliminate bacterial contamination. Cell growth was monitored by the optical density at 610 nm (OD_610_).

### Yeast isolation from the phyllosphere samples

Phyllosphere samples were collected in 2022 from the Kyoto university campus (flowering cherry leaf, satsuma mandarin leaf, ginkgo leaf, loquat leaf, paper mulberry leaf, and satsuki azalea leaf) and a nearby supermarket (apple fruit, broccoli flower bud, grape ’Fujiminori’, ’Nagano purple’, ’Pione’ fruit, and ’Shine Muscat’ fruits, green kiwi fruit, golden kiwi fruit, green onion leaf, lemon fruit, mango fruit, mango peel, natsudaidai orange fruit, natsudaidai orange peel, peach fruit, pineapple fruit, pineapple leaf, pummelo fruit, pummelo peel, and sweet cherry fruit). These samples were cut into small pieces using sterile scissors, and a few pieces were used for cultivation.

Yeast isolation was carried out using an enrichment technique. The samples were inoculated into test tubes containing 5 mL of SM liquid medium and incubated under aerobic conditions. After the first incubation, 50 µL of the suspension was transferred into fresh SM liquid medium for the second round of enrichment. The third enrichment was conducted by inoculating 50 µL from the suspension of the second enrichment into fresh SM liquid medium. Following three consecutive enrichments, each lasting 4 days, a loopful of the suspension was streaked onto SM agar plates and incubated for 3 to 5 days until colonies appeared. Subsequently, 3 to 8 colonies from each plate were picked for identification.

### Identification of yeast species

Yeast genomic DNA was extracted and purified using the MightyPrep reagent (Takara Bio). Briefly, 100 µL of a yeast suspension cultivated on YPD liquid medium was centrifuged at 12,000 rpm for 3 min at 4℃, and the supernatant was removed. The resulting cell pellet was resuspended in 100 µL of MightyPrep reagent. The suspension was then heated at 95℃ for 10 minutes, followed by centrifugation at 12,000 rpm for 5 min at 4℃. The supernatant was transferred to a new tube as the genomic DNA. DNA concentration was measured and adjusted to approximately 200 ng/µL with sterile water.

Using the genomic DNA, D1/D2 domain in the 28S rDNA was PCR-amplified for sequensing analyses using the specific primers NL-1 (5’-GCATATCAATAAGCGGAGGAAAAG-3′) and NL-4 (5’-GGTCCGTGTTTCA AGACGG-3′) (O’donnell, 1993). The internal transcribed spacer (ITS) regions in the 28S rDNA were also PCR-amplified for the representative yeast strains, using the specific primers ITS1 (5’-TCCGTAGGTGAACCTGCGG-3′) and ITS4 (5’-TCCTCCGCTTATTGATATGC-3′) (White et al., 1990). PCR was performed using KOD FX Neo polymerase (Toyobo). The reaction samples with a total volume of 50 µL were prepared by mixing 0.3 µL of diluted genomic DNA, 5.3 µL of PCR-grade water, 12.5 µL of 2× buffer, 5.0 µL of a dNTP mix, and 0.7 µL each of 10 µmol/L forward and reverse primers. The PCR condition was as follows: initial denaturation at 94℃ for 2 min, followed by 30 cycles of denaturation at 98°C for 10 seconds, annealing at 55°C for 30 seconds, and extension at 68°C for 40 seconds.

Subsequently, sequencing analyses of the D1/D2 domain and ITS regions were performed, and the DNA sequences were provided by Eurofin Genomics (Tokyo, Japan). To identify the yeast species, DNA sequences were compared against sequences available in the National Center for Biotechnology Information (NCBI) database. Since the sequences of the D1/D2 domain and ITS regions of *Aureobasidium melanogenum* and *A. pullulans* were not distinguishable, morphological characteristics on the malt extract agar plates were observed for species identification (Gostinčar et al., 2014). Representative yeast strains of this study are listed in Supplemental Table 1.

### Construction of the yeast strains

*C. boidinii* strain TK62 (*ura3*) (Sakai et al., 1991) and *C. boidinii* strain *aod1Δ* (Nakagawa et al., 1999) were transformed with the vector pACT1-Venus, encoding Venus under the control of the *ACT1* promoter (Kawaguchi et al., 2011), and the SK^+^-Zc vector (Kawaguchi et al., 2011), resulting in the zeocin-resistant strain *C. boidinii* strain venus-zeo^R^ and *C. boidinii* strain *aod1Δ* venus-zeo^R^*. P. laurentii* strain PeF4 (this study) was transformed with SK^+^-Zeo^r^ (Yano et al., 2009), yielding the zeocin-resistant strain *P. laurentii* strain zeo^R^. Transformation was conducted using the Fast Yeast Transformation Kit (GE Healthcare, IL, USA), and transformants were selected on agar plates containing 150 µg/mL zeocin.

### Cultivation of *A. thaliana* inoculated with yeasts

*C. boidinii* strain venus-zeo^R^, *C. boidinii* strain *aod1Δ* venus-zeo^R^ and *P. laurentii* strain zeo^R^ were inoculated either under single inoculation or mixed inoculation conditions to compare their colonization abilities on *A. thaliana*. These strains were first grown on YPD liquid medium for 24 h, then transferred to 5 mL of SD liquid medium and cultured for an additional 12-16 h. Cells were collected, washed with sterilized water, and suspended in 0.5 mL sterilized water to prepare a yeast suspension with an OD_610_ of 0.01. For mixed inoculations, 0.25 mL of each yeast suspension was combined in a single tube. *A. thaliana* seeds were surface-sterilized by treatment with 70% ethanol for 1 min, followed by 0.5% (v/v) antiformin containing 0.3% v/v Tween 20 for 15 min, and then washed 4 to 6 times with sterilized water. The sterilized seeds were soaked in 0.5 mL of either the single or mixed yeast suspension for 2 h with gentle shaking at 2 rpm using a Rotator RT-5 (Taitec, Saitama, Japan) at 4℃. Following inoculation, these seeds were grown under aseptic conditions in plastic dishes (approximately 4 cm in height) containing Hoagland agar. Cultivation was carried out in an NK Biotron LH-220 growth chamber (Nippon Medical and Chemical Instruments, Osaka, Japan) at 25°C, 65% humidity, under a 16-h light/8-h dark cycle for 2 weeks.

### Microscopic observation

Microscopic analyses were conducted using the FV3000 confocal microscope (Olympus) to observe *C. boidinii* venus-zeo^R^ cells on *A. thaliana* first leaves. FV3000 is equipped with six solid-state diode lasers (405 nm, 445 nm, 488 nm, 514 nm, 561 nm, and 640 nm) and uses Olympus objectives including PLAPON 60x (1.42 NA, oil-dipped) and UPLSAPO 100x (1.35 NA, Si oil-dipped). High-sensitivity GaAsP detectors are installed in the FV3000, allowing the detection of faint fluorescence signals with high signal-to-noise ratios. Venus fluorescence was obtained with a 488-laser excitation with all diode lasers and LED illumination and a 500-600 nm filter for emission. The transmitted light image was provided using Nomarski difference interference contrast (DIC).

### Colony formation assay

The aerial parts of *A. thaliana* seedlings were excised, weighed, and mixed with 5 mL of sterilized water in a 5 mL tube. The mixture was sonicated using an ultrasonic cleaning device (UT205S, Sharp, Osaka) for 15 min. The resulting aqueous phase was diluted and plated onto SD agar plates supplemented with 150 µg/mL of zeocin. After two days of incubation, colonies appeared on the plates were counted. Fluorescent colonies of *C. boidinii* venus-zeo^R^ were visualized using the FAS-Digi imaging system (NIPPON Genetics).

### Preparation of meythylesterified PGA

Methyl-esterified PGA, both ^13^C-labeled and unlabeled, was prepared, as previously described (Keppler et al., 2008, van Alebeek et al., 2000). Briefly, 2 g of PGA (Sigma-Aldrich, 95 %) were weighed into a 50 mL volumetric flask and supplemented with 15 mL of 0.02 N methanolic sulfuric acid, with the methanol consisting of either 5% ^13^C-labeled methanol (Sigma Aldrich, 99%, 99 atom% ^13^C) or unlabeled methanol (Sigma Aldrich, ≥ 99.8%). The flask was incubated at 30°C for 25 days with shaking at 180 rpm. After incubation, the sample was centrifuged at 4000 rpm for 5 min and the supernatant was discarded. The remaining samples were washed 3 times with 40 mL of propan-2-ol, centrifuging after each step and discarding the supernatant. The washed sample material was left at 50°C in a gas chromatography (GC) oven overnight to allow residual solvent to evaporate. Afterwards, the sample was washed with ultrapure water and then lyophilized for 2 days. The resulting product was ground to a fine powder using a mortar and pestle.

### Definition of stable carbon isotope ratios and δ values

Stable carbon isotope ratios are expressed in the “delta” (δ) notation as the relative difference of the ^13^C/^12^C ratio of a sample compared to the Vienna Peedee Belemnite standard, reported as the δ^13^C value. The unit of delta stable isotope values is given in urey (Ur; 1 mUr = 1 ‰) following the recommendation of (Brand and Coplen, 2012) to align with the International System of Units (SI) guidelines.

### Analysis of methylester content and ^13^C-value of ^13^C-labeled methyl-esterified PGA

The methylester group content was determined using the modified Zeisel method, as previously described (Keppler et al., 2007, Greule et al., 2009). Briefly, 250 µL hydroiodic acid (HI, 57 % (w/v) aqueous solution, Acros; Thermo Fisher Scientific) was added to 30 ± 0.03 mg of ^13^C-labeled or unlabelled methyl-esterified PGA in 1.5 mL crimp-top glass vials. The vials were sealed and heated in an oven at 130°C for 30 min, inducing the reaction between the methylester groups of PGA and hydroiodic acid (HI) to form iodomethane (CH_3_I). After equilibration to room temperature, CH_3_I concentration was analyzed using a gas chromatograph (GC, HP 6890N, Agilent Technologies) equipped with a flame ionization detector (FID) similar to (Li et al., 2012). For the determination of ^13^C-isotopic composition of methyl-esterified PGA, samples were treated with the Zeisel method as described above and an aliquot of headspace (70µL) was injected into a GC (HP 6890N, Agilent Technologies) coupled with a combustion isotope ratio mass spectrometry (C-IRMS, Delta^PLUS^ XL IRMS, ThermoQuest Finnigan; Greule et al., 2009), resulting in a final δ^13^C value of +4069 mUr of ^13^C-labeled methyl-esterified PGA.

### Analysis of ^13^C-CO_2_ formation from ^13^C-labeled PGA

Yeast cells precultured in YPD liquid medium for 24 h were transferred to 10 mL of SD medium at OD_610_ = 0.4 and incubated for 16 h. Subsequently, cells were transferred to 12 mL exetainers (Labco, UK), containing 4 mL of synthetic liquid medium supplemented with 0.5% (w/v) ^13^C-labeled methylesterified PGA (1.8% methylester content, 5% ^13^C-labeled methylester group) or unlabeled methylesterified PGA (1.6% methylester content) at OD_610_ = 5. Cells were grown for 24 h in the exetainers, and the gas in the headspace of the exetainers was collected for analysis. Then, samples were first analyzed for their CO_2_ concentration using a GC coupled with a barrier discharge ionization detector (BID; Shimadzu, Kyoto, Japan) as previously described (Schroll et al., 2024). Subsequently, δ^13^C-CO_2_ values of the samples were analyzed using a GC (HP 6890N, Agilent Technologies) coupled with a Delta^PLUS^ XL IRMS (ThermoQuest Finnigan), similar to the method previously employed (Schroll et al., 2020).

### Cultivation conditions using *Beppu* flasks

Yeasts were co-cultivated using *Beppu* flasks on SP or SD liquid medium. A *Beppu* flask is equipped with a central membrane filter (pore size 0.22 µm) that separates the flask into two compartments. This filter allows the exchange of medium components, such as carbon sources, but prevents the passage of yeast cells. Therefore, yeast strains can share nutrients while their growth is measured separately. After autoclaving, 15 mL of medium was added to each compartment. Strains were pre-cultivated on YPD liquid medium for 24 h, and cells were then transferred into each compartment at an initial OD_610_ of 0.01.

The experimental design for assessing yeast growth is illustrated in Fig. 2B. Cell growth under mono-cultivations was represented by the average OD_610_ values from both sides of the flask. Maximum cell yield was defined as the highest OD_610_ value observed before any decline. The maximum cell yield of *P. laurentii* strain PeF4 co-cultured with each of the *C. boidinii* strains was compared to that of *P. laurentii* strain PeF4 grown in monoculture and expressed as “Fold”.

### Specific extracellular PME activity assays

Yeast cells pre-cultured on YPD liquid medium for 24 h were transferred to 50 mL of SD liquid medium and incubated for 16 h. Subsequently, cells were inoculated into 50 mL of SP or SPM liquid medium to induce PME activity. Cells grown for 8 h or 10 h were harvested by centrifugation, and the supernatants (culture media) were collected as extracellular fractions.

PME activity was measured by titration with 0.01 N NaOH. The reaction mixture consisted of 10 mL of 0.4% pectin in 50 mM acetate buffer (pH 5.0) and 1 mL of extracellular fractions. An aliquot of the same sample was boiled for 10 min and used as a blank control. The reaction mixtures were incubated at 30℃ for 60 min, and the reactions were then stopped by boiling for 10 min. The stopped reaction mixtures were titrated with 0.01 N NaOH to determine the volume needed to reach the same pH as the blank. One unit of PME activity was defined as the amount of enzyme releasing 1 µmol of carboxy groups per min.

### Data availability statement

The sequences of the D1/D2 domain of *Aureobasidium melanogenum* GiL12, *Candida boidinii* SML1, *Cryptococcus* sp. FCL26, *Cystobasidium calyptogenae* PeF7, *Cystobasidium minutum* GSMF2, *Cystobasidium slooffiae* GiL13, *Meyerozyma caribbica* GoKF3, *Naganishia diffluens* ApF1, *Papiliotrema aurea* FCL21, *Papiliotrema laurentii* PeF4, *Pichia kluyveri* GoKF5, *Pseudozyma hubeiensis* SAL1, *Pseudozyma pruni* FCL20, *Rhodosporidiobolus ruineniae* GiL14, *Rhodotorula paludigena* GSMF1, *Rhodotorula toruloides* PeF8, *Sungouiella intermedia* GoKF2 and *Zalaria obscura* GFF1 were deposited in the DDBJ/EMBL/GenBank under accession numbers LC882735, LC882737, LC882739, LC882741, LC882743, LC882745, LC882747, LC882749, LC882751, LC882753, C882755, LC882757, LC882759, LC882761, LC882763, LC882765, LC882767 and LC882769, respectively. Similarly, the sequences of the ITS regions of these yeast strains were deposited under accession numbers LC882736, LC882738, LC882740, LC882742, LC882744, LC882746, LC882748, LC882750, LC882752, LC882754, LC882756, LC882758, LC882760, LC882762, LC882764, LC882766, LC882768 and LC882770, respectively. The original contributions presented in the study are included in the article/Supplementary material; further inquiries can be directed to the corresponding author.

## Author contributions

KShigeta: Conceptualization, Data curation, Formal analysis, Funding acquisition, Investigation, Methodology, Validation, Visualization, Writing – original draft. KShiraishi: Conceptualization, Data curation, Formal analysis, Funding acquisition, Investigation, Methodology, Project administration, Supervision, Validation, Visualization, Writing – original draft, Writing – review & editing. MS: Data curation, Formal analysis, Investigation, Methodology, Validation, Visualization, Writing – review & editing. RL: Data curation, Formal analysis, Investigation, Methodology, Validation, Visualization, Writing – review & editing. FK: Data curation, Formal analysis, Funding acquisition, Methodology, Supervision, Validation, Writing – review & editing. YS: Conceptualization, Formal analysis, Methodology, Supervision, Validation, Writing – review & editing. HY: Conceptualization, Formal analysis, Funding acquisition, Methodology, Supervision, Validation, Writing – review & editing.

## Acknowledgments

We thank S.M. Brosch and J. Kleiner from the Institute of Earth Sciences, Heidelberg University, Germany, for their provision of technical assistance to isotopic labelling analyses. We also thank Dr T. Seike from the Department of Bioscience and Bioinformatics, Kyushu Institute of Technology, and Dr T. Nunoura from the Japan Agency for Marine-Earth Science and Technology for their technical advice about identification of yeast species. KShiraishi appreciates support from the Program for the Development of Next-generation Leading Scientists with Global Insight (L-INSIGHT), sponsored by the Ministry of Education, Culture, Sports, Science and Technology (MEXT), Japan.

## Generative AI statement

The authors declare that no Generative AI was used in the creation of this manuscript.

## Supplementary material

The Supplementary material for this article can be found online.

## Supplementary Information

**Supplemental Fig. 1.**
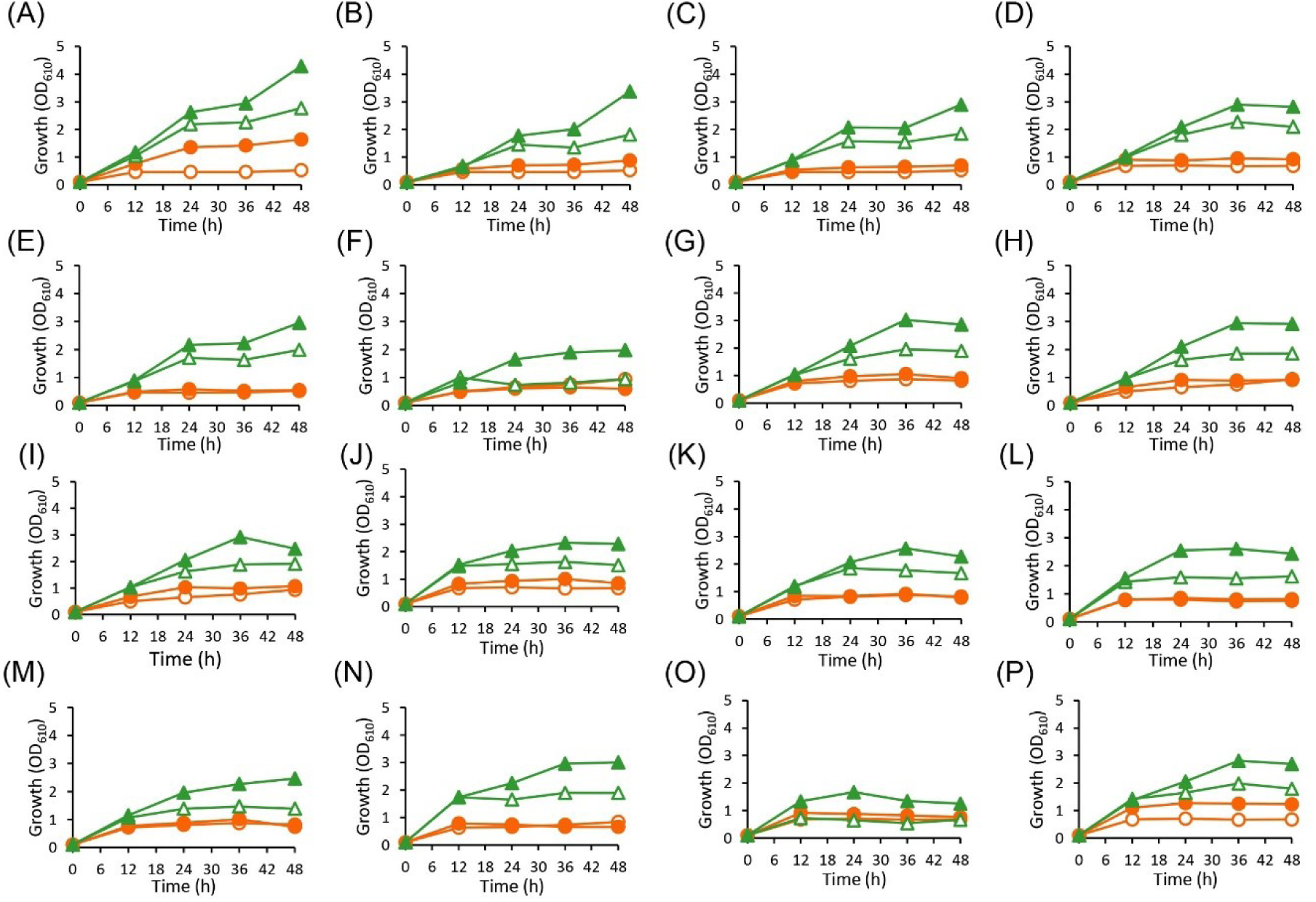
**Cultivation of the other representative yeast strains with *C. boidinii* strain AOU-1 in SP liquid media using *Beppu* flasks.** Growth curve of *C. boidinii* strain AOU-1 and (A) *A. melanogenum* strain GiL12, (B) *Cryptococcus* sp. strain FCL26, (C) *C. calyptogenae* strain PeF7, (D) *C. minutum* strain GSMF2, (E) *C. slooffiae* strain GiL13, (F) *M. caribbica* strain GoKF3, (G) *N. diffluens* strain ApF1, (H) *P. aurea* strain FCL21, (I) *P. kluyveri* strain GoKF5, (J) *P. hubeiensis* strain SAL1, (K) *P. pruni* strain FCL20, (L) *R. ruineniae* strain GiL14, (M) *R. paludigena* strain GSMF1, (N) *R. toruloides* strain PeF8, (O) *S. intermedia* strain GoKF2, and (P) *Z. obscura* strain GFF1. Symbols: (empty circles) *C. boidinii* strain AOU-1 mono-cultivation; (filled circles) *C. boidinii* strain AOU-1 co-cultivation; (empty triangle) each representative yeast strain mono-cultivation; (filled triangle) each representative yeast strain co-cultivation.

**Supplemental Fig. 2.**
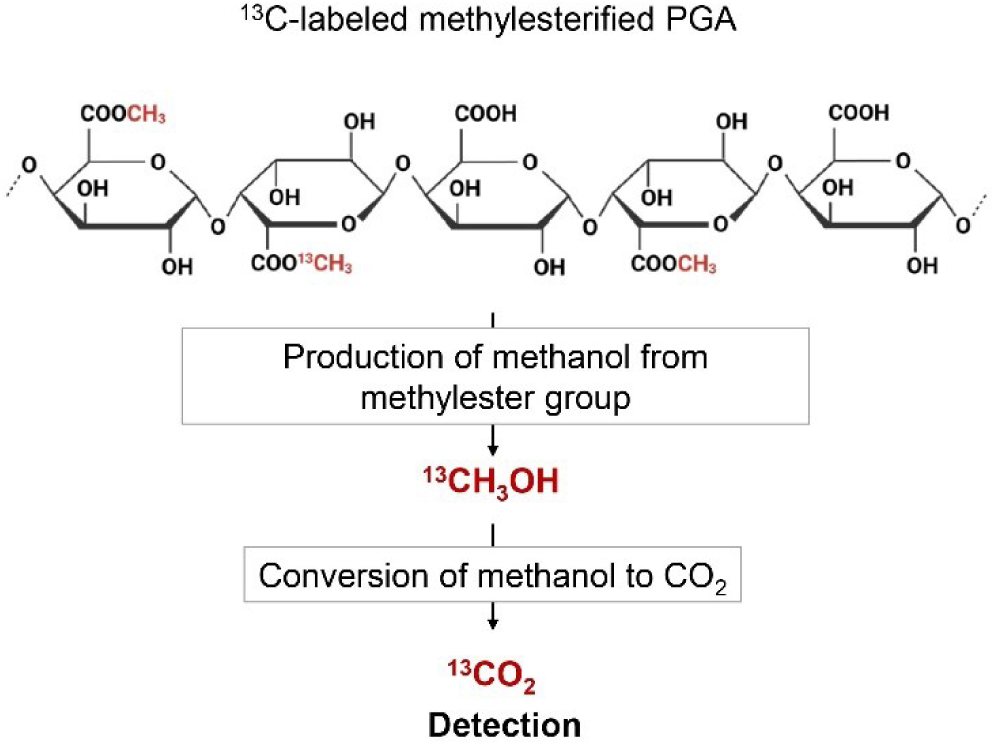
Tracing ^13^C-labeled CO_2_ formation from the methylester group of pectin. Hydrolysis of the ^13^C-labeled methylester group of pectin releases ^13^C-enriched methanol, which is converted to ^13^C-enriched CO_2_, detectable as a final product by gas chromatography-isotope-ratio-mass spectrometry (GC-IRMS).

**Supplemental Fig. 3.**
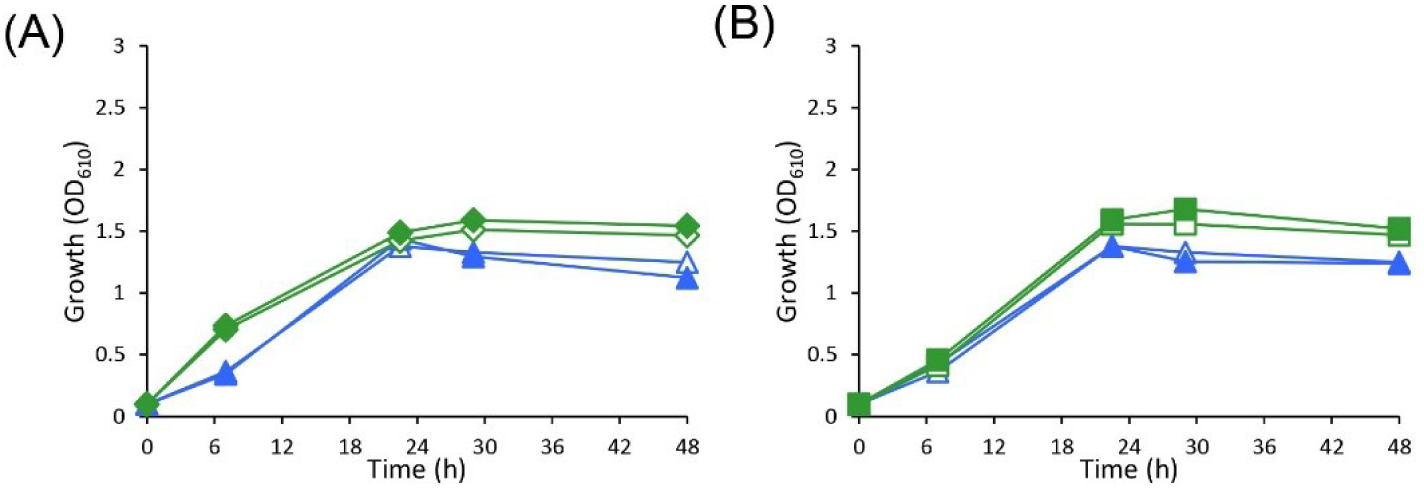
**Cultivation of *P. laurentii* strain PeF4 with *R. ruineniae* strain GiL14 and *R. toruloides* strain PeF8 in SP liquid media using *Beppu* flasks.** Growth curve of *P. laurentii* strain PeF4 and (A) *R. ruineniae* strain GiL14, and (B) *R. toruloides* strain PeF8, during mono- and co-cultivation. Symbols: (empty triangle) *P. laurentii* strain PeF4 mono-cultivation; (filled triangle) *P. laurentii* strain PeF4 co-cultivation; (empty diamonds) *R. ruineniae* strain GiL14 mono-cultivation; (filled diamonds) *R. ruineniae* strain GiL14 co-cultivation;.(empty squares) *R. toruloides* strain PeF8 mono-cultivation; (filled squares) *R. toruloides* strain PeF8 co-cultivation.

**Supplemental Fig. 4.**
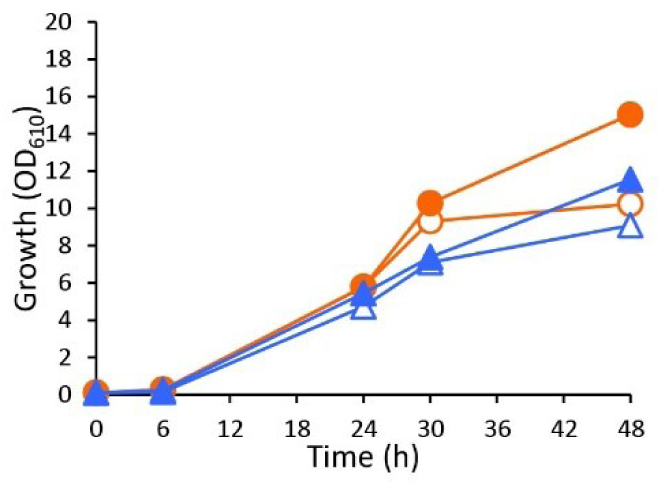
**Cultivation of *P. laurentii* strain PeF4 with *C. boidinii* strain AOU-1 in SD liquid media using *Beppu* flasks.** Growth curve of *C. boidinii* strain AOU-1 and *P. laurentii* strain PeF4 during mono- and co-cultivation. Symbols: (empty circles) *C. boidinii* strain AOU-1 mono-cultivation; (filled circles) *C. boidinii* strain AOU-1 co-cultivation; (empty triangle) *P. laurentii* strain PeF4 mono-cultivation; (filled triangle) *P. laurentii* strain PeF4 co-cultivation.

**Supplemental Fig. 5.**
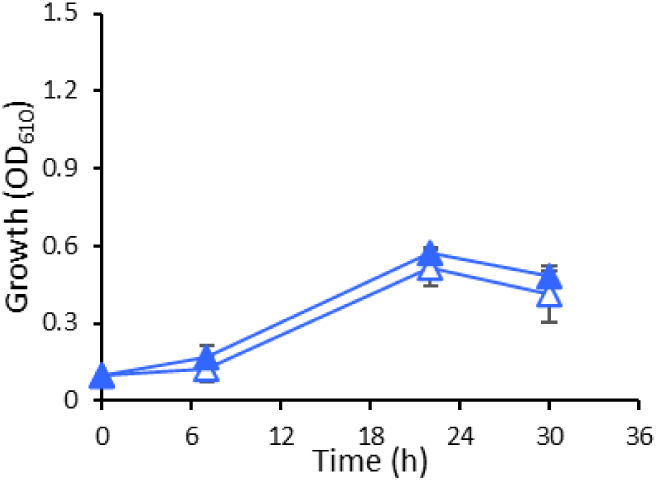
**Growth curve *P. laurentii* strain PeF4 in SP and SPM liquid media.** Symbols: (empty triangles) *P. laurentii* strain PeF4 in SP medium; (filled triangles) *P. laurentii* strain PeF4 in SPM liquid medium.

**Supplemental Fig. 6.**
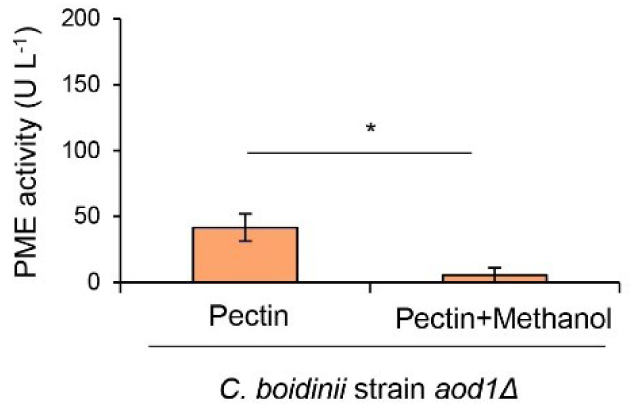
**Specific extracellular PME activity of *C. boidinii* strain *aod1Δ* in SP and SPM liquid media.** The data are presented as the mean±standard error (SE; n = 6). Significant differences are marked by *(*p* < 0.05; Student’s *t*-test).

**Supplemental Table 1:**
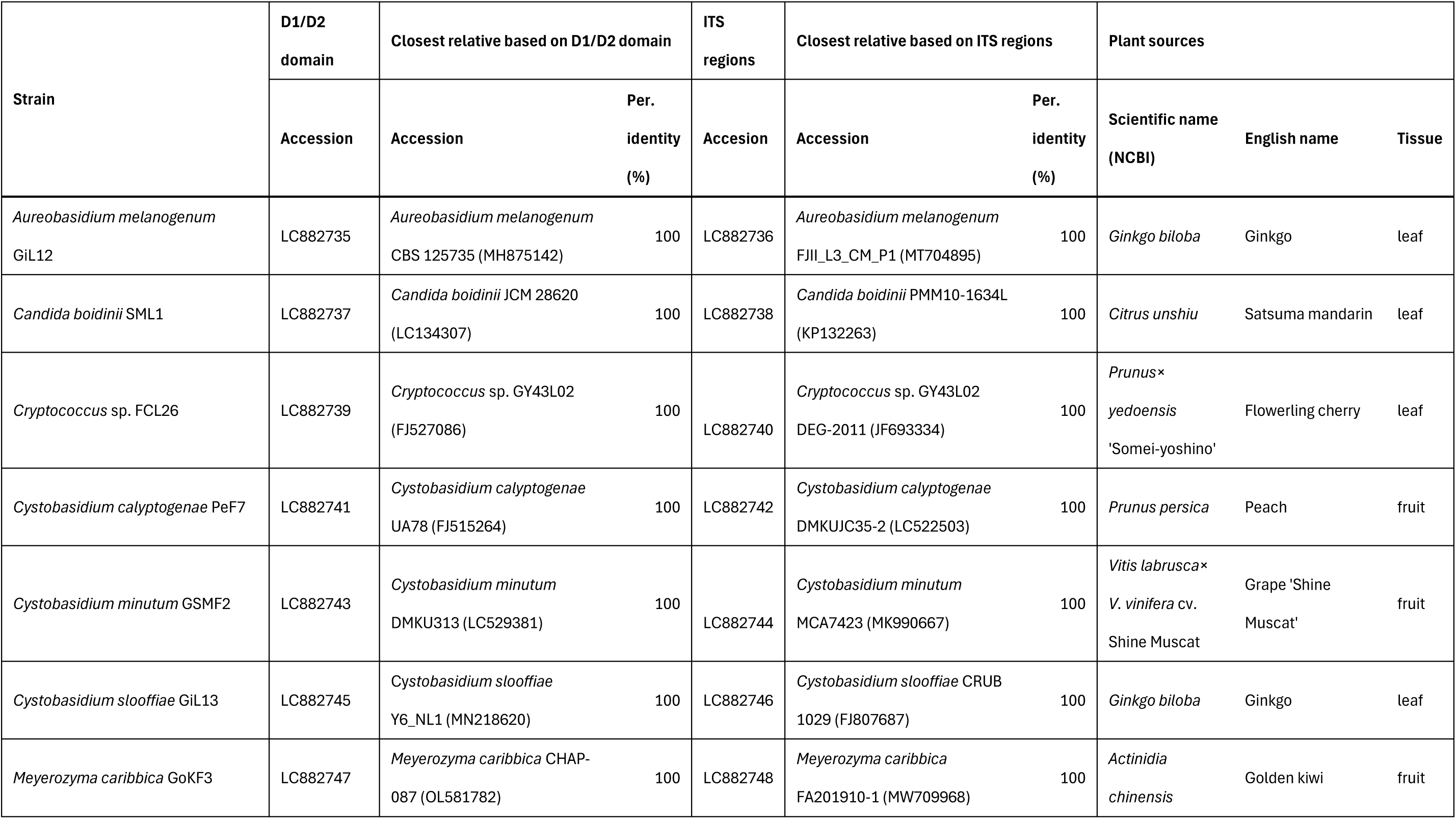
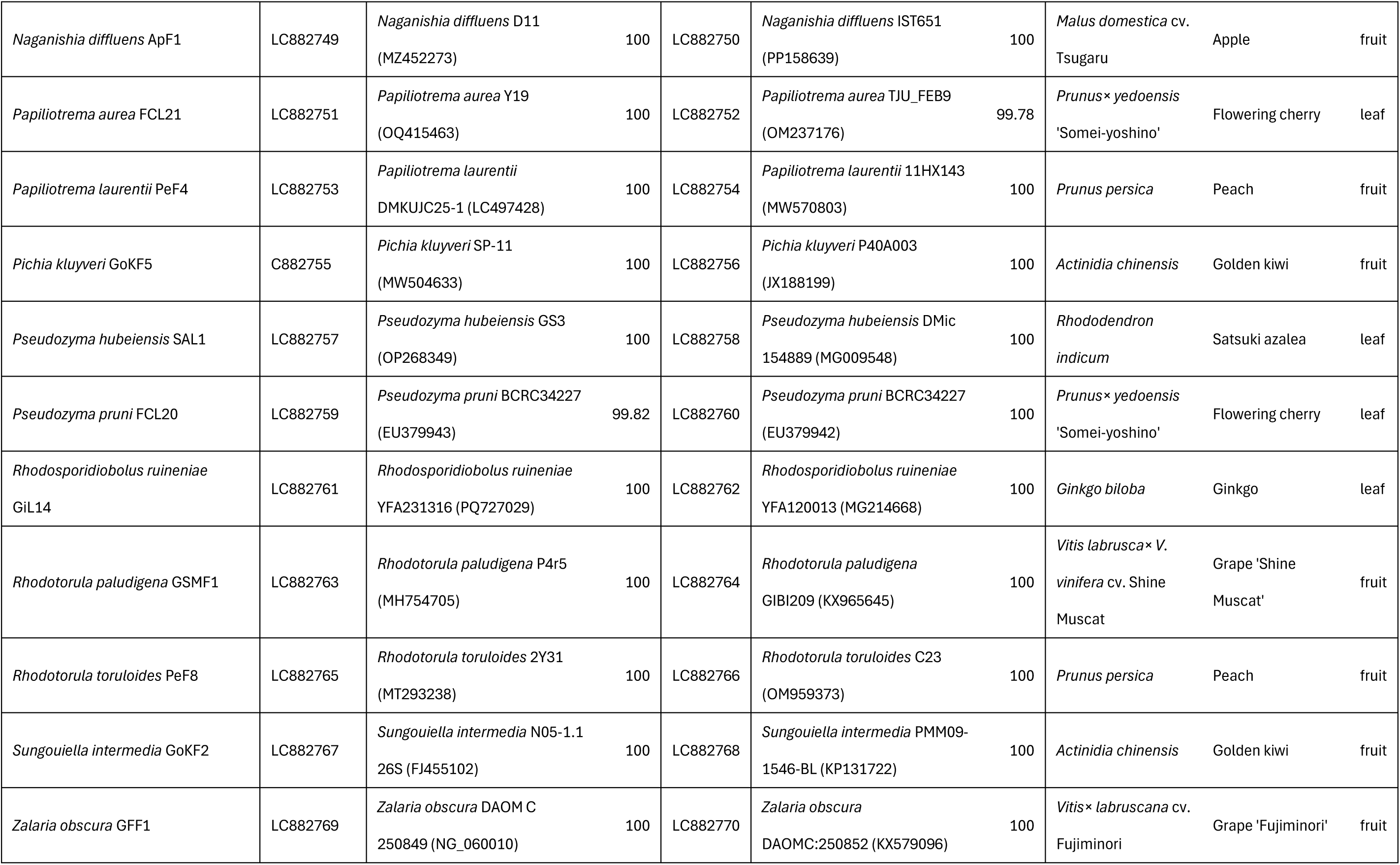
The representative yeast strains isolated and identified from the phyllosphere.

| Strain | D1/D2 domain | Closest relative based on D1/D2 domain |  | ITS regions | Closest relative based on ITS regions |  | Plant sources |  |  |
| --- | --- | --- | --- | --- | --- | --- | --- | --- | --- |
|  | Accession | Accession | Per. identity (%) | Accession | Accession | Per. identity (%) | Scientific name (NCBI) | English name | Tissue |
| *Aureobasidium melanogenum* GIL12 | LC882735 | *Aureobasidium melanogenum* CBS 125735 (MH875142) | 100 | LC882736 | *Aureobasidium melanogenum* FJII_L3_CM_P1 (MT704895) | 100 | *Ginkgo biloba* | Ginkgo | leaf |
| *Candida boidinii* SML1 | LC882737 | *Candida boidinii* JCM 28620 (LC134307) | 100 | LC882738 | *Candida boidinii* PMM10-1634L (KP132263) | 100 | *Citrus unshiu* | Satsuma mandarin | leaf |
| *Cryptococcus* sp. FCL26 | LC882739 | *Cryptococcus* sp. GY43L02 (FJ527086) | 100 | LC882740 | *Cryptococcus* sp. GY43L02 DEG-2011 (JF693334) | 100 | *Prunus*×<br>*yedoensis*<br>'Somei-yoshino' | Flowerling cherry | leaf |
| *Cystobasidium calyptogenae* PeF7 | LC882741 | *Cystobasidium calyptogenae* UA78 (FJ515264) | 100 | LC882742 | *Cystobasidium calyptogenae* DMKUJC35-2 (LC522503) | 100 | *Prunus persica* | Peach | fruit |
| *Cystobasidium minutum* GSMF2 | LC882743 | *Cystobasidium minutum* DMKU313 (LC529381) | 100 | LC882744 | *Cystobasidium minutum* MCA7423 (MK990667) | 100 | *Vitis labrusca*×<br>*V. vinifera* cv.<br>Shine Muscat | Grape 'Shine Muscat' | fruit |
| *Cystobasidium slooffiae* GIL13 | LC882745 | *Cystobasidium slooffiae* Y6_NL1 (MN218620) | 100 | LC882746 | *Cystobasidium slooffiae* CRUB 1029 (FJ807687) | 100 | *Ginkgo biloba* | Ginkgo | leaf |
| *Meyerozyma caribbica* GoKF3 | LC882747 | *Meyerozyma caribbica* CHAP-087 (OL581782) | 100 | LC882748 | *Meyerozyma caribbica* FA201910-1 (MW709968) | 100 | *Actinidia chinensis* | Golden kiwi | fruit |
| *Naganishia diffluens* ApF1 | LC882749 | *Naganishia diffluens* D11<br>(MZ452273) | 100 | LC882750 | *Naganishia diffluens* IST651<br>(PP158639) | 100 | *Malus domestica* cv.<br>Tsugaru | Apple | fruit |
| *Papiliotrema aurea* FCL21 | LC882751 | *Papiliotrema aurea* Y19<br>(OQ415463) | 100 | LC882752 | *Papiliotrema aurea* TJU_FEB9<br>(OM237176) | 99.78 | *Prunus*× *yedoensis*<br>'Somei-yoshino' | Flowering cherry | leaf |
| *Papiliotrema laurentii* PeF4 | LC882753 | *Papiliotrema laurentii*<br>DMKUJC25-1 (LC497428) | 100 | LC882754 | *Papiliotrema laurentii* 11HX143<br>(MW570803) | 100 | *Prunus persica* | Peach | fruit |
| *Pichia kluyveri* GoKF5 | C882755 | *Pichia kluyveri* SP-11<br>(MW504633) | 100 | LC882756 | *Pichia kluyveri* P40A003<br>(JX188199) | 100 | *Actinidia chinensis* | Golden kiwi | fruit |
| *Pseudozyma hubeiensis* SAL1 | LC882757 | *Pseudozyma hubeiensis* GS3<br>(OP268349) | 100 | LC882758 | *Pseudozyma hubeiensis* DMic<br>154889 (MG009548) | 100 | *Rhododendron*<br>*indicum* | Satsuki azalea | leaf |
| *Pseudozyma pruni* FCL20 | LC882759 | *Pseudozyma pruni* BCRC34227<br>(EU379943) | 99.82 | LC882760 | *Pseudozyma pruni* BCRC34227<br>(EU379942) | 100 | *Prunus*× *yedoensis*<br>'Somei-yoshino' | Flowering cherry | leaf |
| *Rhodospiridiobolus ruineniae*<br>GIL14 | LC882761 | *Rhodospiridiobolus ruineniae*<br>YFA231316 (PQ727029) | 100 | LC882762 | *Rhodospiridiobolus ruineniae*<br>YFA120013 (MG214668) | 100 | *Ginkgo biloba* | Ginkgo | leaf |
| *Rhodotorula paludigena* GSMF1 | LC882763 | *Rhodotorula paludigena* P4r5<br>(MH754705) | 100 | LC882764 | *Rhodotorula paludigena*<br>GIBI209 (KX965645) | 100 | *Vitis labrusca*× *V.*<br>*vinifera* cv. Shine<br>Muscat | Grape 'Shine<br>Muscat' | fruit |
| *Rhodotorula toruloides* PeF8 | LC882765 | *Rhodotorula toruloides* 2Y31<br>(MT293238) | 100 | LC882766 | *Rhodotorula toruloides* C23<br>(OM959373) | 100 | *Prunus persica* | Peach | fruit |
| *Sungouiella intermedia* GoKF2 | LC882767 | *Sungouiella intermedia* N05-1.1<br>26S (FJ455102) | 100 | LC882768 | *Sungouiella intermedia* PMM09-1546-BL (KP131722) | 100 | *Actinidia chinensis* | Golden kiwi | fruit |
| *Zalaria obscura* GFF1 | LC882769 | *Zalaria obscura* DAOM C<br>250849 (NG_060010) | 100 | LC882770 | *Zalaria obscura*<br>DAOMC:250852 (KX579096) | 100 | *Vitis*× *labruscana* cv.<br>Fujiminori | Grape 'Fujiminori' | fruit |

